# Transcriptional Profiles of Antidepressant Resistance Across the Corticolimbic Pathway of Chronically Stressed Mice

**DOI:** 10.1101/2025.03.17.643727

**Authors:** Trevonn Gyles, Eric M. Parise, Molly S. Estill, Lyonna F. Parise, Caleb J. Browne, Li Shen, Eric J. Nestler, Angélica Torres-Berrío

**Author notes:** These authors contributed equally to this work (Trevonn Gyles, Eric M. Parise).

## Abstract

Treatment-resistant depression (TRD), defined by unsuccessful response to multiple antidepressants, affects approximately one-third of individuals with major depressive disorder, yet its underlying molecular mechanisms remain poorly understood. Here, we developed a preclinical model of TRD in which mice exposed to chronic social defeat stress were sequentially treated with fluoxetine (FLX) and ketamine (KET), allowing behavioral stratification into antidepressant responsive and non-responsive mice. RNA sequencing of the nucleus accumbens (NAc) and prefrontal cortex (PFC) revealed transcriptional signatures associated with treatment outcomes. Prior exposure to FLX exerted a priming effect that facilitated molecular and behavioral responsiveness to KET in a subset of animals in both the NAc and PFC. However, this priming effect was absent in non-responders, despite identical treatment regimes, suggesting a transcriptional divergence underlying differential outcomes. Gene co-expression network analysis identified modules enriched for differentially expressed genes unique to stress-susceptible and FLX-KET nonresponsive mice, as well as modules overlapping with both stress susceptibility and antidepressant resistance. These findings suggest that failed antidepressant treatment can shape the brain’s molecular landscape in a way that influences subsequent treatment outcomes, and that resistance arises not simply from treatment failure but from an absence of adaptive molecular priming. This work provides insight into the gene networks contributing to antidepressant non-response and highlights a mechanistic framework for modeling TRD in preclinical systems. By identifying molecular correlates of sequential pharmacological resistance, our findings may inform the development of novel therapeutic strategies for individuals with TRD.

## INTRODUCTION

Unsuccessful response to multiple antidepressant treatments is a key signature of treatment-resistant depression (TRD), a serious subtype of major depressive disorder (MDD) that affects nearly 30% of patients undergoing pharmacotherapy [1–5]. MDD is highly heterogeneous, with symptoms that vary according to severity and longitudinal trajectories [6] and an increasing incidence as a direct consequence of the COVID-19 pandemic [7]; however, the lack of response to treatment despite adequate dose, duration, and adherence remains a significant challenge for clinical practice [1–5], highlighting the urgent need to develop novel pharmacological approaches with better therapeutic outcomes [7,8]. A limitation to achieving this goal is the limited understanding of the neurobiology mediating successful versus unsuccessful responses to currently available antidepressants.

Conventional antidepressants have not evolved over the last decades and continue to rely on targeting monoaminergic neurotransmission, such as selective serotonin reuptake inhibitors, or tricyclic antidepressants [8,9]. These agents are the first line choice for pharmacotherapy and are typically selected through trial-and-error given the lack of reliable predictors of antidepressant efficacy [8,10,11]. Moreover, virtually all classes of monoamine-based drugs require several weeks to display symptom improvement [8–12], and nearly half of the patients require multiple rounds of treatment [8–12]. Recently, Ketamine (KET), an antagonist of the glutamate N-methyl-D-aspartate (NMDA) receptor, among other actions, has been shown to induce a rapid and lasting antidepressant response in TRD patients [1,13–16]. Yet, KET has not been approved as an initial choice to treat MDD [15].

Animal models are indispensable for investigating the biological mechanisms of antidepressant response. Yet most preclinical studies examine a single antidepressant regimen in drug-naïve animals, often using behavioral assays that detect acute responses but fail to model the delayed effects seen in patients [17–19]. Therefore, more translationally relevant models are needed to capture the progressive and heterogeneous nature of TRD.

Our group previously used the chronic social defeat stress (CSDS) paradigm to examine transcriptional correlates of antidepressant efficacy in mice [20]. Specifically, Bagot et al. [20] used CSDS to compare the transcriptional profiles associated with successful versus unsuccessful responses to either the tricyclic antidepressant imipramine or KET across key corticolimbic regions, including the nucleus accumbens (NAc), prefrontal cortex (PFC), hippocampus, and amygdala [21–24]. Overall, this work demonstrated that successful response to either antidepressant resembled both pro-resilient and anti-susceptible transcriptional signatures in mice [20]. By contrast, unsuccessful response was associated with pro-susceptible gene expression patterns [20,25]. A question that remains is whether distinct transcriptional profiles are observed in mice that persistently fail to respond to multiple courses of antidepressants and whether these profiles could serve to uncover molecular mechanisms underlying TRD.

Here, we developed a preclinical TRD model in which CSDS-susceptible mice were first treated with chronic fluoxetine (FLX) and subsequently administered a single dose of KET. We performed RNA-sequencing in both the NAc and PFC, core components of limbic circuitry that mediate mood, executive function, and stress regulation, and are highly sensitive to antidepressant-induced plasticity [21–24]. We selected FLX because it is among the first-line pharmacotherapies for treating MDD [1,11], and it is well-tolerated in mice when administered in drinking water [26,27], providing a more naturalistic and translational route of administration as compared to intraperitoneal injections [28]. We found that prior failure of FLX response primes a successful treatment with KET. We also identified a subset of mice that persistently failed to respond to FLX and KET and uncovered gene signatures associated with treatment resistance to both drugs.

## MATERIALS AND METHODS

### Animals

All the experimental procedures were approved by the Institutional Animal Care and Use Committee at the Icahn School of Medicine at Mount Sinai. Male C57BL/6J wild-type mice (postnatal day 75±15, Jackson Laboratory) were used as experimental mice and maintained on a 12-hour light/dark cycle (lights on at 07:00) with food and water ad libitum. CD-1 male breeders (≥3 months old, Charles River) were used as aggressors. Experimental mice were group-housed before stress exposure and single-housed thereafter. Behavioral analyses were conducted by experimenters blinded to treatment groups.

### Chronic Social Defeat Stress (CSDS)

CSDS was performed as in [29]. Briefly, each experimental mouse was exposed to 5-minute of physical attacks by a CD-1 aggressor. After the session, experimental and CD-1 mice were housed overnight in a 2-compartment hamster cage and separated by a transparent divider with holes to provide sensory, but not physical, contact. The procedure was repeated during 10 consecutive days, in which experimental mice faced a new aggressor. Control C57BL/6J mice were housed in 2-compartment cages with a different cage-mate every day. The CSDS protocol was conducted between 11:00 and 14:00.

### Social Interaction Test (SIT)

The SIT was assessed twenty-four hours after the last CSDS exposure. The SIT consisted of 2 sessions (2.5-minute, each), in which experimental mice explored a squared-arena (44 x 44 cm) in the absence or presence of a novel CD-1 aggressor (social target). In the first session, an empty wired-mesh enclosure (10 x 6.5x 42 cm) was located against one of the walls of the arena to assess baseline exploration. In the second session, an unfamiliar social target was placed inside the wired-mesh enclosure. The area surrounding the enclosure was designated as the social interaction zone (SIZ) (14 x 9 cm). The time (in seconds) in the SIZ was measured during both sessions. The social interaction ratio (time in SIZ with social target/time in SIZ without social target) was calculated to classify mice as susceptible (ratio 1) and resilient (ratio≥1) [29]. This simple measure correlates strongly with numerous other behavioral outcomes, including cognitive flexibility [30–32].

### Fluoxetine (FLX) Treatment

FLX (Spectrum Chemicals) was dissolved in regular drinking water at a 160mg/L concentration and delivered ad libitum for twenty-eight days through Black Light-Safe conical tubes capped with rubber stoppers. FLX concentration and route of administration were selected based on [26,27]. Water-treated susceptible and control mice received regular drinking water in the same type of tubes. FLX and water consumption were measured and replaced every third day. Twenty-four hours after the final FLX treatment day, all mice were assessed on a second SIT (SIT2). Mice with >25% social interaction improvement from SIT1 to SIT2 were classified as FLX-responders; others were classified as non-responders.

### Ketamine (KET) Treatment

KET (Vedco) was dissolved in saline and administered as a single injection (10mg/kg, i.p.) to FLX non-responders and a subset of saline-treated susceptible mice twenty-four hours after the SIT2 [20]. A third SIT (SIT3) was assessed twenty-four hours post-injection.

### Tissue Dissection

NAc and PFC tissue punches were collected twenty-four hours after the SIT3. All experimental mice were euthanized by rapid decapitation. Brains were removed, cooled with ice-cold PBS and sliced on a pre-defined brain matrix. Unilateral PFC punches (12-gauge) and bilateral NAc punches (14-gauge) were collected from 1mm coronal sections starting on approximately plate 15 of the mouse brain atlas [33], and frozen immediately on dry ice.

### RNA Isolation, Library Preparation and RNA-Sequencing

Total RNA from NAc and PFC tissue was isolated with the RNeasy Mini Kit (Qiagen). All RNA samples were determined to have values ≥1.8 with Nanodrop (Thermo Fisher), and RNA integrity ≥8 with the RNA 6000 Nano Bioanalyzer Assay (Agilent). Libraries were prepared using a 50ng purified RNA concentration with the TruSeq Stranded Total RNA Kit with ribosomal RNA depletion (Illumina). Libraries were size selected and purified using AMPure XP beads (Beckman Coulter), and concentrations were quantified with the High-Sensitivity DNA Bioanalyzer Assay (Agilent). Libraries were sequenced at a 40 million paired-end reads on an Illumina NovaSeq 6000 System (GENEWIZ/AZENTA).

### RNA-Seq Data Analysis

Quality control was performed using FASTQC software (https://github.com/shenlab-sinai/NGS-Data-Charmer). Raw reads underwent adapter trimming and were mapped to the mouse genome (mm10) using HISAT [34]. Count matrices were generated using the feature Counts function of the Subreads program (https://subread.sourceforge.net). Differential gene expression (DEGs) was analyzed using the DESeq2 package [35]. Genes with low counts (<10 counts in <5 samples) were excluded from downstream analysis. The following cutoffs were applied to identify DEGs: Log2(fold change) > |0.20|, nominal p-value of <0.05, and false discovery rate <0.1.

### Pattern Analysis for Differential Expression

To cluster gene groups with similar expression trajectories across experimental conditions, we used the degPatterns function from the DEGreport R package [36]. Only DEGs passing statistical significance thresholds were included as inputs. The degPatterns applies hierarchical clustering based on pairwise correlations among normalized expression values and automatically determines the number of gene clusters by optimizing within-and between-cluster variability. This approach allowed us to identify co-regulated gene groups and visualize condition-specific transcriptional patterns.

### Multiscale Embedded Gene Co-Expression Network Analysis (MEGENA)

MEGENA was used to construct gene co-expression networks in NAc and PFC samples separately [37,38]. Networks were built from variance-stabilized counts across 37 samples, yielding 61,769 (NAc) and 62,343 (PFC) edges. Modules with <50 genes were excluded. Hub genes were defined by node strength and degree (FDR <0.05), and enrichment statistics were calculated using the GeneOverlap package (https://github.com/shenlab-sinai/GeneOverlap). Top 25% hub genes were visualized using the igraph and ggraph R packages with the Fruchterman– Reingold layout for gene connectivity [37,39]. This force-directed layout emphasizes proximity of highly co-expressed gene nodes and reveals module structure.

### Gene Ontology (GO) Enrichment Analysis

GO biological processes were identified using the enrichGO clusterProfiler v2023 [40]. Enriched terms were visualized using tidyverse and ggplot2.

### Quantification and Statistical Analysis

Data are presented as mean ± SEM. Significance was defined as p < 0.05. Parametric or nonparametric tests were used based on normality and variance assumptions. Analyses were performed in GraphPad Prism 6.0. Group differences were evaluated using two-tailed t-tests or ANOVAs with Tukey’s or Sidak’s post hoc tests. Outliers were identified via the ROUT method. No statistical methods were used to predetermine sample sizes.

## RESULTS

### Behavioral Responses to Consecutive Courses of Antidepressant Treatments

To model aspects of TRD, adult male mice were subjected to CSDS and stratified as susceptible or resilient based on the social interaction test (SIT) [29,30,32,41,42]. Mice exhibiting reduced social interaction in the initial SIT (SIT1) were classified as susceptible and received either fluoxetine (FLX; 160 mg/L in drinking water; SUS-FLX) or regular drinking water (SUS-W) **(Figure 1A-C)**. Following the 28-day treatment, control (CON), SUS-W, and SUS-FLX mice were assessed in a second SIT (SIT2). Among FLX-treated mice, ∼65% showed increased social interaction and were classified as responders (FLX-RESP), while ∼35% remained socially avoidant (FLX-NR) **(Figure 1D–E)**, consistent with response variability observed in both clinical and preclinical studies [1,2,20,41–43]. Water consumption was comparable across groups, ruling out dosing differences as a confound **(Figure 1F; Figure S1)**.

**Figure 1.**
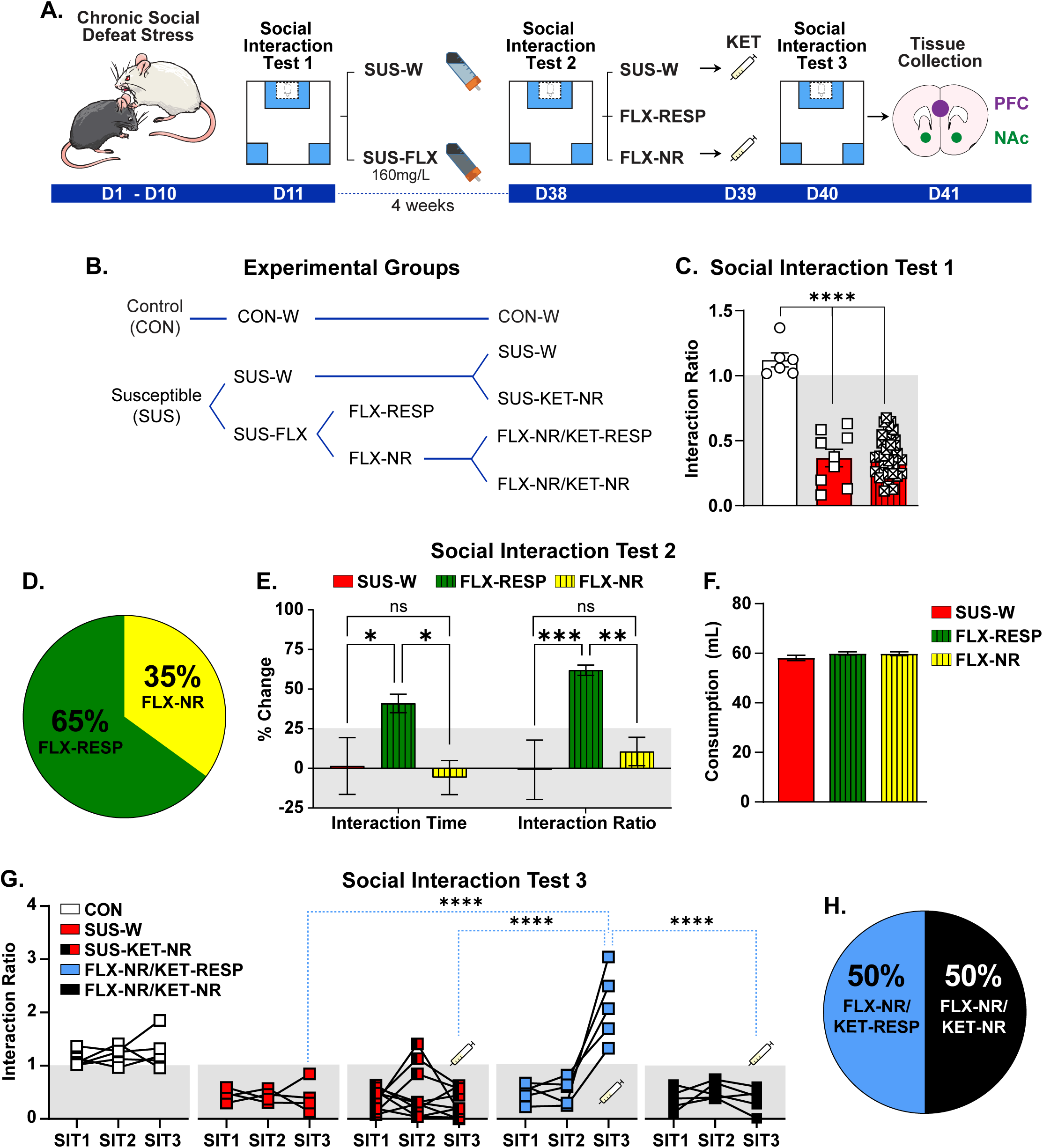
Successful versus unsuccessful response to consecutive courses of antidepressants in susceptible mice after CSDS. (**A**) Schematic and timeline of the CSDS experiment and fluoxetine (FLX) and ketamine (KET) administration. (**B**) Experimental groups: Control (CON), Water-treated susceptible (SUS-W), KET-non-responders (SUS-KET-NR), FLX-treated susceptible (SUS-FLX), FLX-responders (FLX-RESP), FLX-non-responders (FLX-NR), FLX-non-responders/KET-responders (FLX-NR/KET-RESP), and FLX-non-responders/KET-non-responders (FLX-NR/KET-NR). (**C**) Social interaction test 1 (SIT1). Interaction ratio prior to FLX treatment: One-way ANOVA: F(2,45)= 54.24; p<0.0001. Tukey’s test: SUS-W and SUS-FLX different from CON, ****p<0.0001. (**D**) Percentage of responder (FLX-RESP) and non-responder (FLX-NR) mice to FLX treatment. (**E**) Percentage of change from social interaction test 1 to test 2. Left: Change in time of interaction with social target: One-way ANOVA: F(2,40)= 11.33; p<0.001. Tukey’s test: Increased interaction time in FLX-RESP compared to SUS-W (*p<0.05) and FLX-NR (*p<0.05). Right: Change in Interaction ratio: One-way ANOVA: F(2,40)= 19.32; p<0.0001. Tukey’s test: Increased ratio in FLX-RESP compared to SUS-W (***p<0.001) and FLX-NR (**p<0.01). **(F)** Average of total water consumption during FLX treatment. One-way ANOVA: F(2,30)= 54.24; p=0.26. (**G**) Social interaction ratio across antidepressant treatments. Two-way ANOVA: Group: F(3,48)= 15.61; p<0.0001; Social interaction test: F(2,48)= 6.79; p<0.01; Group by Social interaction test interaction: F(6,48)=12.51; p<0.0001. Tukey’s test: Increased interaction ratio in FLX-NR/KET-RESP compared to SIT 3 in SUS-W, SUS-KET-NR, and FLX-NR/KET-NR, ****p<0.0001, and compared to SIT1 and SIT2 in FLX-NR/KET-RESP, ***p<0.0001. (**H**). Percentage of responders (FLX-NR/KET-RESP) and non-responders (FLX-NR/KET-NR) to KET treatment in mice that failed to respond to FLX. Data: mean ± SEM.

To assess the effects of sequential antidepressant treatments, FLX-NR mice and half of the SUS-W group received a single intraperitoneal injection of ketamine (KET; 10 mg/kg) and were re-evaluated in a third SIT (SIT3) 24 hours later **(Figure 1A–B)**. Approximately 50% of FLX-NR mice exhibited improved social interaction (FLX-NR/KET-RESP), while the remainder continued to display social avoidance (FLX-NR/KET-NR) **(Figure 1G–H)**. Notably, none of the SUS-W mice responded to a single injection of KET (SUS-KET-NR), suggesting that prior FLX exposure, even if behaviorally ineffective, may facilitate subsequent KET efficacy **(Figure S2-3)**. This observation may also reflect methodological differences from prior studies, since we administered KET four weeks after CSDS, compared to two weeks in earlier work (e.g., Bagot et al. [20]), raising the possibility that the timing of KET delivery influences its behavioral efficacy. Because we did not observe a clear behavioral responder group in the SUS-KET condition and our timing diverged from previously validated protocols, we excluded this group from primary transcriptomic comparisons. Nevertheless, despite the absence of a behavioral effect, SUS-KET mice exhibited a distinct transcriptional signature in both the NAc and PFC when compared to SUS-W and FLX-RESP groups **(Figure S2-3)**. These findings suggest that even in the absence of overt behavioral improvement, KET engages molecular pathways that may have therapeutic relevance and underscore the importance of continued investigation into its potential as a first-line antidepressant. Together, our results demonstrate that stress-susceptible mice vary in their responses to consecutive antidepressant administration.

### Transcriptional Signatures of Single versus Sequential Antidepressant Treatments

To characterize the transcriptional effects of antidepressant treatments in stress-susceptible mice, we compared gene expression in the NAc and PFC across four experimental groups: SUS-W, FLX-RESP, FLX-NR/KET-RESP, and FLX-NR/KET-NR, relative to stress-naive CON. This approach enabled us to compare the full impact of single-agent and sequential treatments on the transcriptional architecture of two key brain regions implicated in MDD and treatment response [11,15,16,20,44–47].

Differential gene expression (DEGs) relative to CON revealed a striking reversal of the stress-induced transcriptional profile in SUS-W mice across all treatment groups, including those that failed to show behavioral recovery **(Figure 2A)**. Although SUS-W mice exhibited robust upregulation and downregulation of gene expression in both brain regions, all three antidepressant-treated groups displayed an opposing transcriptional direction, suggesting that drug exposure induces a strong molecular response that can counteract stress-related alterations even in the absence of overt behavioral rescue. This pattern was consistent in both the NAc and PFC.

**Figure 2.**
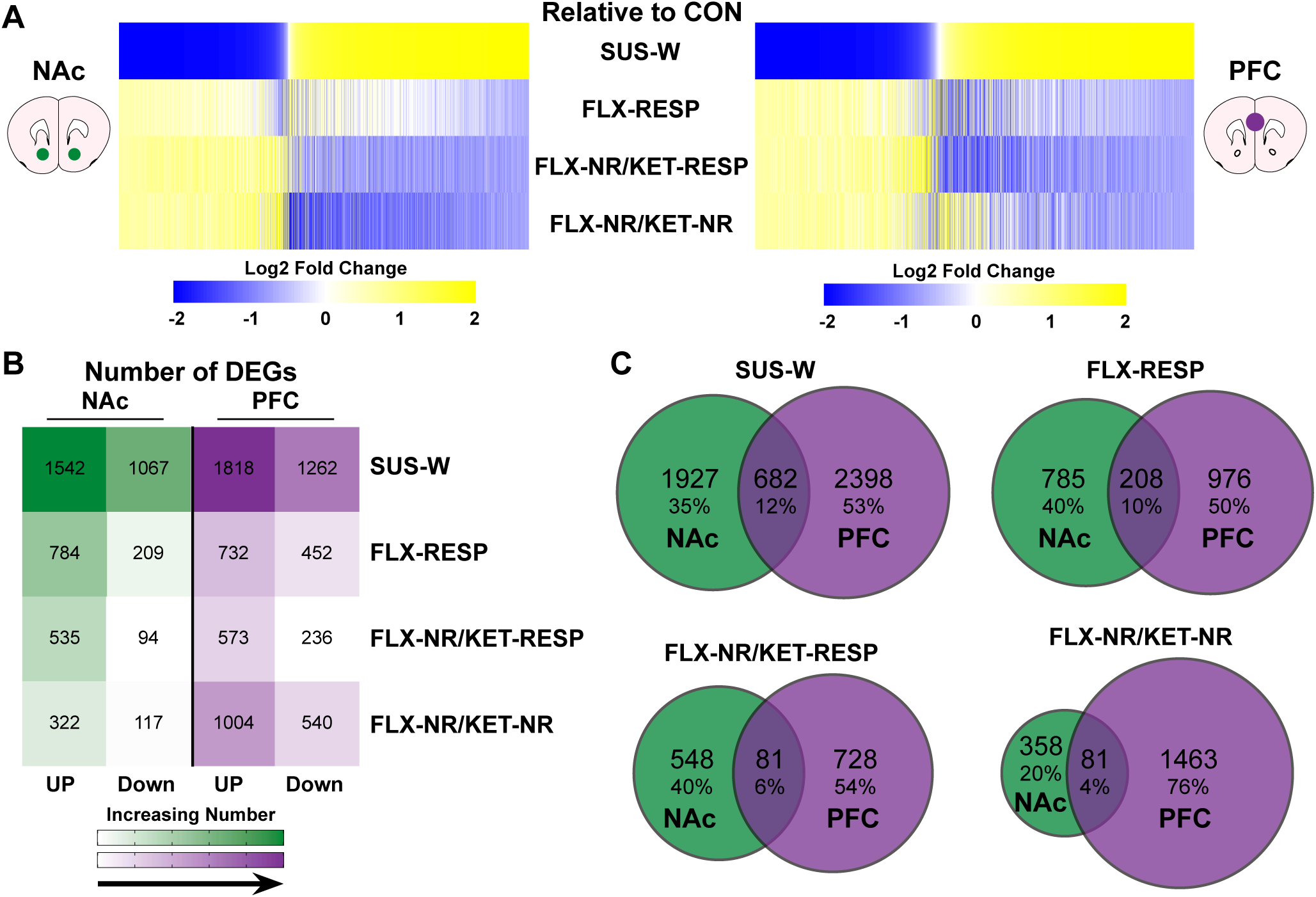
Transcriptional effect of antidepressant treatment in NAc and PFC relative to stress-naive controls. **(A)** Heatmaps displaying union DEGs (relative to CON) across NAc (left) and PFC (right) for each group: SUS-W, FLX-RESP, FLX-NR/KET-RESP, and FLX-NR/KET-NR. Genes are organized based on the SUS-W vs. CON comparison. DEG criteria: Log2(fold change) > |0.20|, nominal p-value of <0.05, and false discovery rate <0.1. Blue and yellow represent downregulation and upregulation, respectively. **(B)** Number of DEGs (up and down) in NAc and PFC for each group relative to CON. Color scale indicates increasing gene count. **(C)** Venn diagrams showing the number of DEGs unique or shared across brain regions for each experimental group. The highest overlap was observed in SUS-W, with decreasing overlap across FLX-RESP, FLX-NR/KET-RESP, and FLX-NR/KET-NR groups, indicating reduced regional concordance with increasing pharmacological intervention.

We observed the largest number of DEGs in SUS-W mice, followed by FLX-RESP, FLX-NR/KET-RESP, and FLX-NR/KET-NR mice **(Figure 2B)**. Notably, the transcriptional response was more pronounced in the PFC than in the NAc across all groups, and all drug-treated groups exhibited fewer DEGs than SUS-W mice. This transcriptional attenuation suggests that both single and sequential antidepressant treatments may dampen the transcriptional “tone” of stress susceptibility, even in cases where behavioral improvement is not observed.

Finally, we then generated Venn diagrams to assess the overlap of DEGs for each group and across brain regions **(Figure 2C)**. The SUS-W condition showed the greatest overlap across regions, followed by FLX-RESP, FLX-NR/KET-RESP, and FLX-NR/KET-NR groups. This progressive reduction in shared DEGs may reflect increasing regional specificity of treatment-induced transcriptional changes. Together, these data demonstrate that antidepressant exposure, whether behaviorally effective or not, alters gene expression in a manner that is both region-specific and divergent from the stress-induced state, and that the PFC shows greater transcriptional responsivity to these interventions than the NAc.

### Distinct Molecular Signatures of Single versus Sequential Antidepressant Treatment Relative to Stress Susceptibility

To identify antidepressant-induced transcriptional signatures and disentangle treatment effects from baseline susceptibility, we compared gene expression across FLX-RESP, FLX-NR/KET-RESP, and FLX-NR/KET-NR groups relative to SUS-W. Interestingly, we found that both FLX-NR/KET-RESP and FLX-NR/KET-NR mice exhibited transcriptomic profiles that were broadly inverse to those observed in FLX-RESP mice in NAc and PFC **(Figure 3A)**. This mirrored reversal across both brain regions suggests that KET exerts distinct transcriptional effects when added following FLX, regardless of behavioral response, potentially signifying a pharmacological shift in underlying mechanisms.

**Figure 3.**
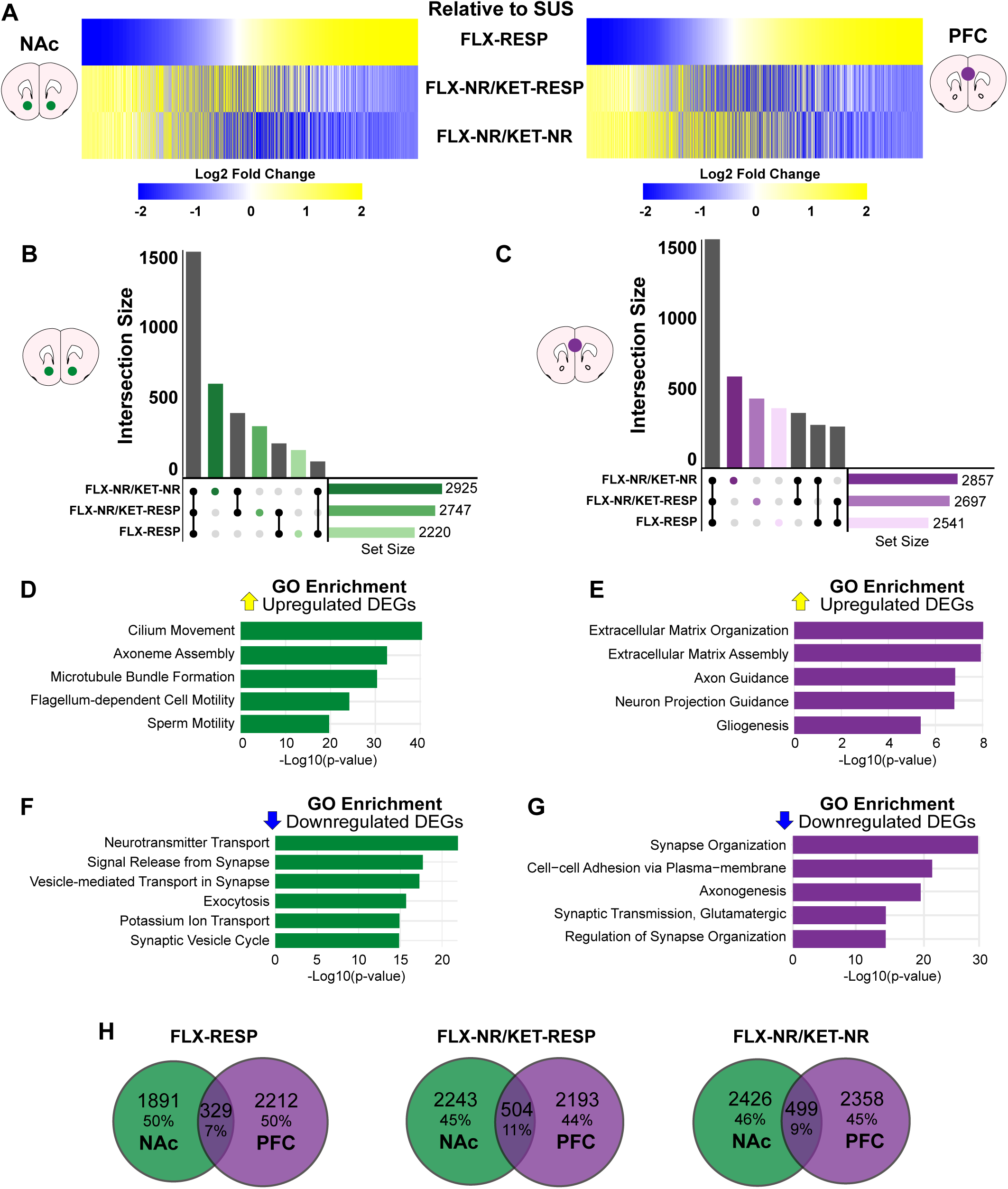
Sequential antidepressant treatment induces distinct transcriptional signatures relative to stress-susceptible mice. **(A)** Heatmaps show DEGs in the NAc (left) and PFC (right) for FLX-RESP, FLX-NR/KET-RESP, and FLX-NR/KET-NR groups, using SUS-W as the reference. Genes are organized based on the FLX-RESP vs. SUS-W comparison. **(B-C)** UpSet plots display DEG set sizes (bottom right) and intersections (bar height) for each group relative to SUS-W in the NAc **(B)** and PFC **(C)**. Most DEGs are shared across groups, with additional group-specific DEGs, especially in sequentially treated mice. **(D-G)** GO enrichment analysis for upregulated (top) and downregulated (bottom) DEGs in NAc **(D-E)** and PFC **(F-G)**, highlighting processes related to microtubule formation, synaptic signaling, and extracellular matrix remodeling. **(H)** Venn diagrams show DEG overlap between NAc and PFC for each group. While functional categories are shared, most DEGs are region-specific, with only modest cross-regional overlap (7–11%).

To quantify DEG distribution across conditions, we generated UpSet plots showing both the total number of DEGs and their intersection across groups **(Figure 3B–C)**. More than 1,500 DEGs were shared across all three treatment groups in both NAc and PFC, with additional genes uniquely altered in each group. The highest number of unique DEGs was observed in FLX-NR/KET-NR mice, followed by FLX-NR/KET-RESP and FLX-RESP. Notably, the two groups receiving sequential antidepressants exhibited greater transcriptional similarity to each other than to the FLX-RESP group, supporting that sequential treatment induces a distinct molecular program not observed with FLX alone.

We then performed Gene Ontology (GO) enrichment analysis on up-and downregulated DEGs in each brain region **(Figure 3D–E)**. In the NAc, upregulated genes were enriched for terms related to microtubule bundle formation, or axoneme assembly, while downregulated genes were associated with neurotransmitter transport, synaptic vesicle cycling, and exocytosis. In the PFC, upregulated genes were associated with extracellular matrix organization and axon guidance, whereas downregulated genes were enriched for synapse organization and glutamatergic signaling pathways. These results suggest that antidepressant treatment engages molecular processes involved in structural remodeling and synaptic regulation, with region-specific emphasis.

Despite similarities in functional enrichment across regions, majority of DEGs were region-specific **(Figure 3F)**. Indeed, the DEGs overlap between NAc and PFC was quite modest, with 7% for FLX-RESP, 11% for FLX-NR/KET-RESP, and 9% for FLX-NR/KET-NR, highlighting brain region-specific regulation of gene expression in response to antidepressants. Nonetheless, the modest gene overlap may represent core components of a cross-regional antidepressant signature.

### FLX Non-Responders Exhibit Distinct, Non-Inverse, Transcriptional Signatures Following KET Treatment

To further isolate the molecular signatures associated with lack of antidepressant response, we compared FLX-NR/KET-RESP and FLX-NR/KET-NR mice relative to FLX-RESP mice. This strategy isolates the transcriptional effects of KET treatment in FLX-treated animals and contrasts responders with non-responders. Union heatmaps revealed that neither FLX-NR/KET-RESP nor FLX-NR/KET-NR mice displayed simple inverse expression patterns in the NAc and PFC **(Figure 4A)**. This suggests that lack of behavioral response may not result from opposing molecular programs, but rather from diminished priming effects induced by prior FLX exposure. Canonical pathway analysis revealed convergent activation of several neurobiologically relevant pathways, including synaptogenesis, CREB signaling, serotonin receptor signaling, and extracellular matrix (ECM) remodeling **(Figure 4B–C)**. Notably, ECM organization, a pathway implicated in synaptic plasticity and antidepressant efficacy, was among the top upregulated pathways in both NAc and PFC [48]. While this functional overlap suggests shared pathway engagement, the underlying genes appeared partially distinct between responders and non-responders.

**Figure 4.**
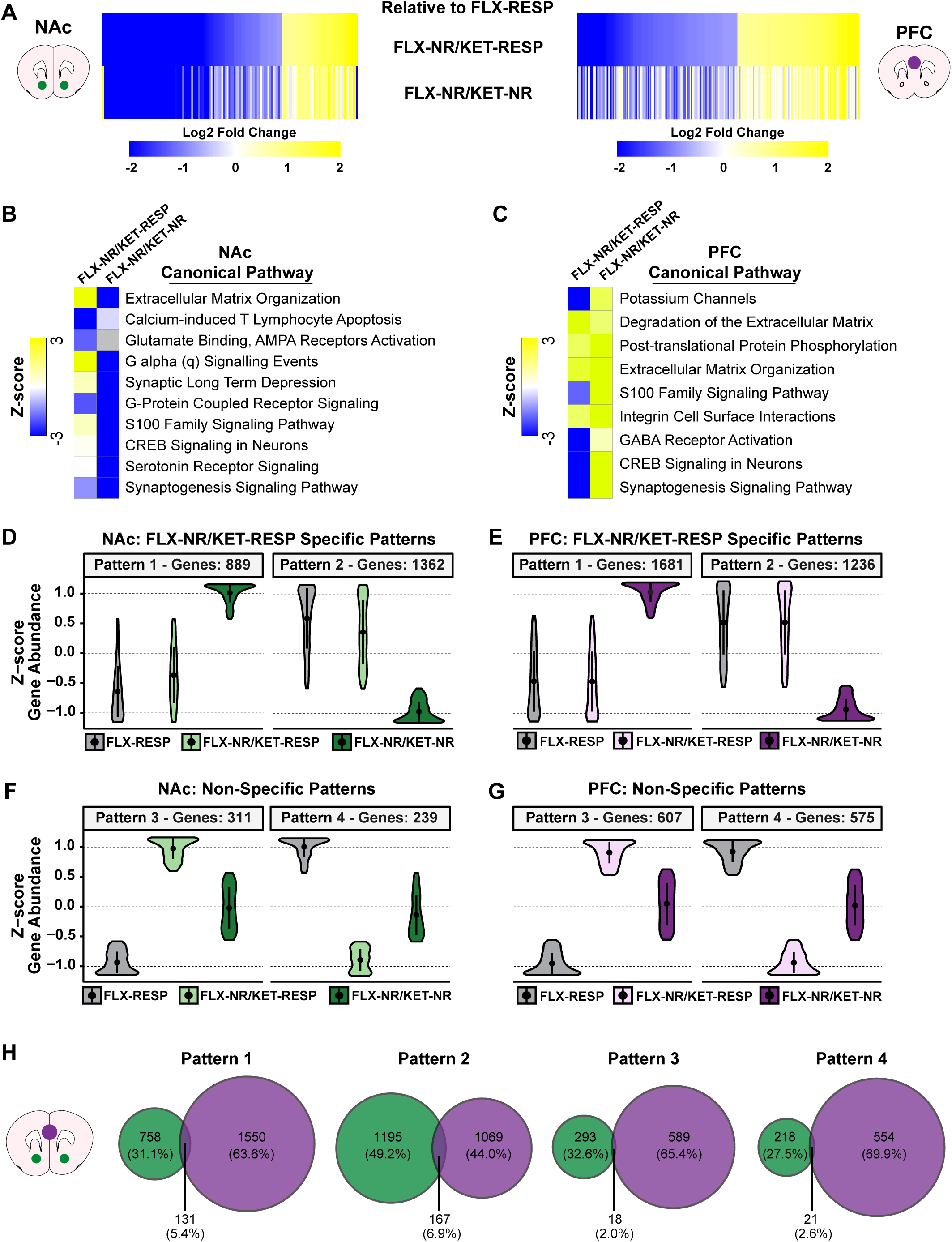
FLX-NR/KET-NR mice exhibit distinct transcriptional and pathway signatures compared to FLX responders. **(A)** Heatmaps showing DEGs in NAc and PFC of FLX-NR/KET-RESP and FLX-NR/KET-NR groups relative to FLX-RESP. Genes are clustered based on the FLX-RESP vs. SUS-W comparison. **(B–C)** Ingenuity Pathway Analysis (IPA) showing top canonical pathways for each group in NAc **(B)** and PFC **(C)**, ranked by Z-score. Pathways include extracellular matrix remodeling, CREB signaling, serotonin receptor signaling, and synaptogenesis. **(D-E)** Violin plots show z-scored gene abundance for FLX-NR/KET-NR-specific clusters (Clusters 1–2) in NAc and PFC. These clusters include genes uniquely regulated in FLX-NR/KET-NR mice. **(F-G)** Non-specific clusters (Clusters 3–4) include genes similarly regulated across all groups. **(H)** Venn diagrams display DEG overlap between NAc and PFC for each cluster. Overlap is modest (5–7%), suggesting region-specific regulation of both unique and shared transcriptional programs.

We also identified two FLX-NR/KET-NR-specific patterns (patterns 1 and 2), comprising genes that were uniquely upregulated or downregulated in this group compared to both FLX-RESP and FLX-NR/KET-RESP **(Figure 4D-E)**. In contrast, non-specific patterns (patterns 3 and 4) included genes similarly regulated across all groups **(Figure 4F-G)**. These FLX-NR/KET-NR-specific clusters likely capture molecular processes associated with persistent lack of response. Despite broad pathway similarity, the overlap of gene clusters between NAc and PFC was very low, with the highest overlap at only ∼5% **(Figure 4H)**. These results emphasize the regional specificity of gene regulation in response to antidepressants and highlight patterns 1 and 2 as molecular candidates for treatment resistance. Together, these findings suggest that lack of response to sequential FLX and KET treatment is not driven by a mirror image of the response signature, but by a qualitatively distinct transcriptional program particularly in a subset of genes that appear uniquely dysregulated in non-responders.

### Gene Co-Expression Networks Reveal Region-Specific Signatures of Antidepressant Response and Resistance

To examine whether transcriptional changes associated with antidepressant response or non-response are organized within coordinated gene networks, we performed Multiscale Embedded Gene Co-expression Network Analysis (MEGENA) separately in the NAc and PFC. We then overlaid DEGs from SUS-W, FLX-RESP, FLX-NR/KET-RESP, and FLX-NR/KET-NR groups (relative to control) onto the respective MEGENA networks to visualize enrichment patterns across modules. In the NAc, two modules showed robust DEGs enrichment for SUS-W and FLX-NR/KET-NR mice but were not enriched in either treatment responder group **(Figure 5A)**. This pattern suggests that these modules are associated with stress susceptibility and are persistently dysregulated in treatment-resistant animals. In contrast, successful response to treatment, whether via a single or sequential antidepressant regimen, may normalize or bypass these transcriptionally vulnerable modules.

**Figure 5.**
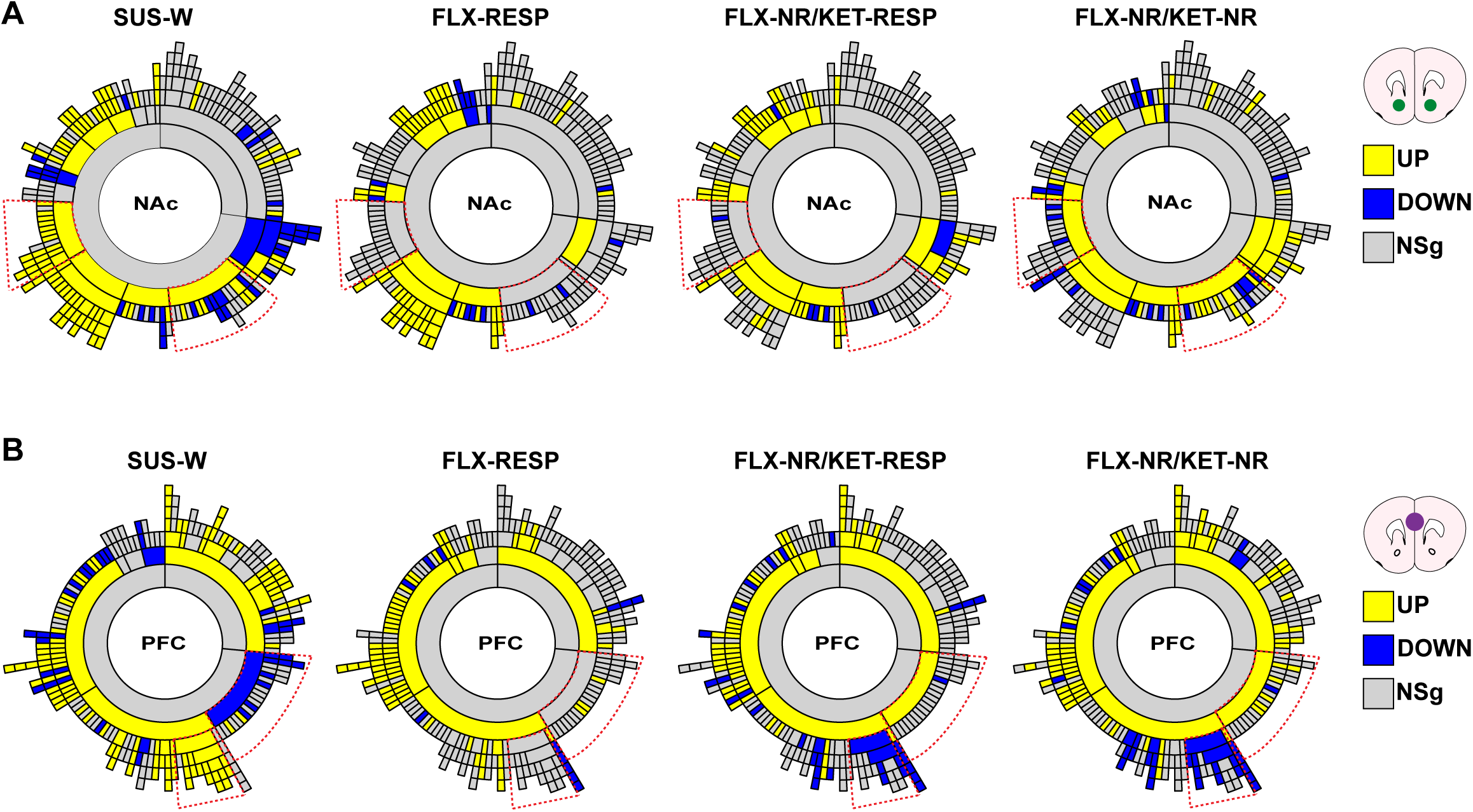
Gene co-expression networks reveal stress-and treatment-specific module enrichment in NAc and PFC. **(A)** Sunburst plots showing MEGENA-derived network modules in the NAc with DEG enrichment overlay for each group (relative to control): SUS-W, FLX-RESP, FLX-NR/KET-RESP, and FLX-NR/KET-NR. Two modules are enriched for upregulated genes in SUS-W and FLX-NR/KET-NR mice only. **(B)** Equivalent plots for the PFC show two stress-enriched modules that are reversed in both FLX-NR/KET-RESP and FLX-NR/KET-NR mice, suggesting shared PFC transcriptional effects of KET. Direction of enrichment is color-coded: yellow (upregulated), blue (downregulated), gray (no significant enrichment). Red dotted arcs indicate modules of interest. Modules enriched in FLX-NR/KET-NR but not in responders may underly treatment resistance in the NAc.

In the PFC, two modules were also enriched for DEGs in SUS-W mice **(Figure 5B)**. However, unlike in the NAc, both FLX-NR/KET-RESP and FLX-NR/KET-NR groups exhibited a reversal of the direction of gene regulation within these modules. This pattern was not observed in FLX-RESP mice and suggests that the PFC may exhibit a broader, KET-associated molecular response that is partially independent of the behavioral outcomes. The fact that both sequential treatment groups showed similar enrichment in the PFC, regardless of response, points to a potential role for this region in mediating core molecular effects of KET.

Finally, to explore the molecular architecture of lack of antidepressant response, we visualized the network structure of a key treatment-resistant modules identified in the NAc MEGENA output (**Figure S4**). This module was strongly enriched for DEGs in the FLX-NR/KET-NR group and contained two densely interconnected subnetworks centered around **SNAP25** and **SYNGR1**, genes with established roles in synaptic transmission and psychiatric illness [49,50]. Their centrality within the treatment-resistant module suggests that altered vesicle trafficking and presynaptic signaling may underlie persistent lack of antidepressant response.

Collectively, these findings reveal a region–specific architecture of antidepressant response with the PFC exhibiting transcriptional plasticity in response to treatment regardless of the behavioral outcome, whereas the NAc appears to harbor persistent, stress-associated gene networks that are selectively active in non-responders.

## DISCUSSION

Antidepressant resistance remains a major clinical challenge, with a substantial proportion of patients failing to respond to first-line pharmacotherapies despite adequate dose and duration [1–5]. While fast-acting agents like KET have emerged as promising alternatives [13,15,46,47,51], the neurobiological mechanisms governing responses versus non-responses—especially following sequential treatments—are not well understood [1–5]. Using the CSDS rodent model, we investigated how prior exposure to FLX influences behavioral and transcriptional outcomes following KET treatment. By stratifying mice into responders and non-responders to both single and combinatorial antidepressant regimens, we identified region–specific gene expression and network-level signatures that distinguish successful treatment from persistent resistance.

Our behavioral data revealed that ∼65% of FLX-treated susceptible mice showed improved social interaction, while a substantial subset remained non-responsive, recapitulating pre-clinical response variability [20,52,53]. Likewise, our results are consistent with human studies reporting diverse degrees of FLX efficacy in MDD-treated individuals [1–5]. Our results also showed that mice treated with KET following an unsuccessful response to FLX separated into responders (50%) and non-responders (50%). Strikingly, we observed that none of the water-treated susceptible mice that received a single injection of KET showed an antidepressant-like effect to this drug. While previous evidence has shown that a single KET injection reverses social avoidance in nearly 50% of susceptible mice to CSDS [20,53], our new findings suggest that the rate of KET success may be lower if the administration of this drug occurs longer after CSDS exposure (e.g., 2 weeks versus 4 weeks); therefore, aspects such as timing or treatment duration can impact the behavioral response to KET in mice. Moreover, this data further indicates that previous FLX administration, even if it fails to alleviate stress-induced behavioral abnormalities, may promote successful response to subsequent KET treatment. In this context, it would be critical to assess whether alternative antidepressant compounds [54,55] or the combination of other types of conventional pharmacotherapies [20] produce parallel behavioral effects to those induced by FLX.

Robust evidence shows that while FLX and KET initially act via serotonin reuptake and glutamate receptors [13,15,46,47,51,56], both converge on common intracellular cascades linked to neuronal plasticity [28,57,58]. Consistent with this, our transcriptional data revealed partial overlap in gene regulation between FLX responders and FLX non-responders who subsequently responded to KET, suggesting that FLX primes molecular programs that facilitate KET efficacy. Thus, sequential antidepressants may involve overlapping signaling pathways but elicit divergent outcomes based on prior priming and gene responsiveness.

MEGENA network analysis revealed that unsuccessful treatment preserved stress-related transcriptional modules, particularly in the NAc, where two modules were enriched in SUS-W and FLX-NR/KET-NR mice but not in responders. Network visualization of a resistance-enriched NAc module identified two prominent hub genes: **SNAP-25** and **SYNGR1,** with known roles in synaptic transmission [49,50]. SNAP-25, a key SNARE complex protein, is essential for neurotransmitter release and has been implicated in depression [49,59], while SYNGR1, a synaptic vesicle protein, has previously emerged as a hub in CSDS-susceptible mice [60]. These hubs may coordinate maladaptive connectivity in resistant animals, reinforcing the idea that treatment failure involves active transcriptional reprogramming within synaptic networks.

Despite its strengths, this study has limitations. It focused exclusively on male mice, limiting generalizability given known sex differences in depression prevalence [61,62], stress response, and antidepressant effects [48,63–65]. Future work must include both sexes to define sex-specific TRD mechanisms. Additionally, behavioral outcomes were assessed using the SIT, which reliably captures stress susceptibility [29] and correlates with broader behavioral phenotypes [29,30,32,42], but does not fully capture the multidimensional nature of TRD. Further testing should incorporate reward-based and cognitive assessments to align with human symptom domains [66,67]. Moreover, the CSDS model offers an opportunity to integrate machine learning approaches for phenotyping in naturalistic settings [31,53,63,68–71], increasing translational relevance.

In summary, this study is the first to define transcriptional changes associated with successful versus unsuccessful responses to sequential antidepressant treatments in chronically stressed mice. By uncovering distinct gene networks across the NAc and PFC, our findings lay the foundation for mechanistic studies aimed at overcoming treatment resistance. These insights will help shape future approaches to personalize antidepressant therapy and target molecular substrates of TRD.

## ACKNOWLEDGMENTS

AUTHOR CONTRIBUTIONS

A.T.B. E.M.P., and E.J.N. designed and conceptualized the study. A.T.B., E.M.P., T.G., M.S.E., and L.F.P., conducted experiment and analyzed the data. A.T.B., T.G., and E.J.N wrote the manuscript. All authors discussed, commented on and edited the paper.

## FUNDING

This work was supposed by grants from the National Institute of Mental Health (R01MH129306 to EJN), the Hope for Depression Research Foundation (to EJN), the Robin Chemers Neustein Award and the Friedman Brain Institute Innovation Award (to ATB) and the Brain & Behavior Research Foundation award (30609 to EPM). ATB is a Charles H. Hood Foundation Scholar and is supported by the Massachusetts General Hospital Lurie Center for Autism, Department of Pediatrics, and Center for Academic Development and Enrichment.

## COMPETING INTERESTS

The authors have nothing to disclose.

## DATA AND MATERIALS AVAILABILITY

All data needed to evaluate the conclusions in the paper are present in the paper and/or the Supplementary Materials. All RNAseq data reported in this study will be deposited publicly in the Gene Expression Omnibus upon manuscript acceptance. Other supporting scripts/code used in this study are available from the corresponding author upon request.

## Notes

### Competing Interest Statement

The authors have declared no competing interest.

### Summary of Updates

We included an additional dataset and modified the text and the main figures.

