## Supplementary figures and images for "Transcriptional Profiles of Antidepressant Resistance Across the Corticolimbic Pathway of Chronically Stressed Mice"

### Figure S1.

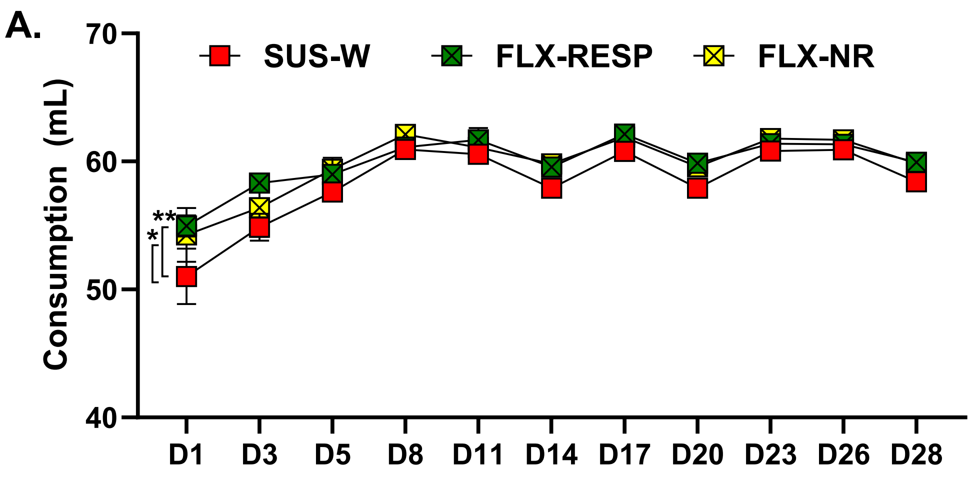

### Figure S2.

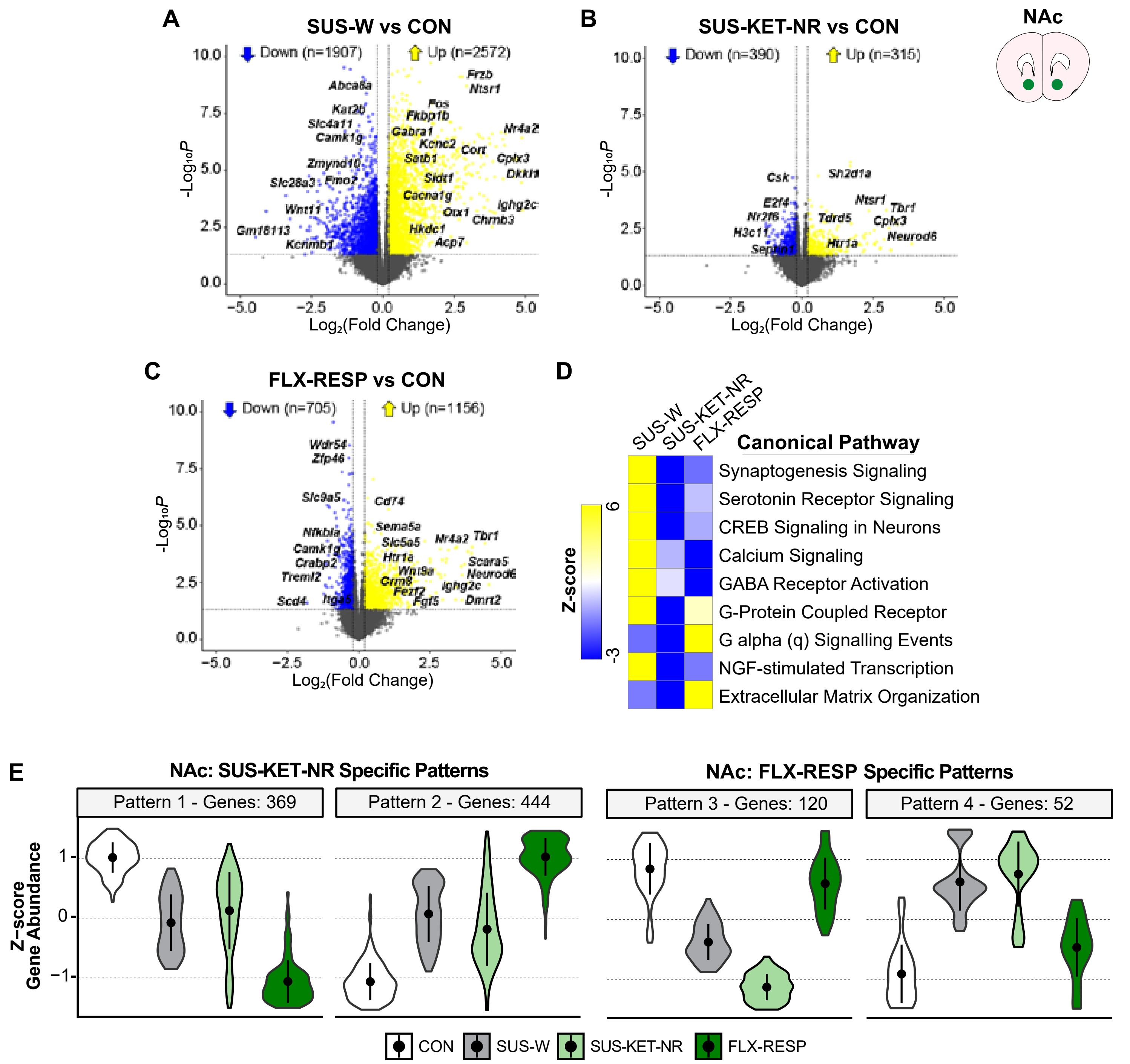

### Figure S3.

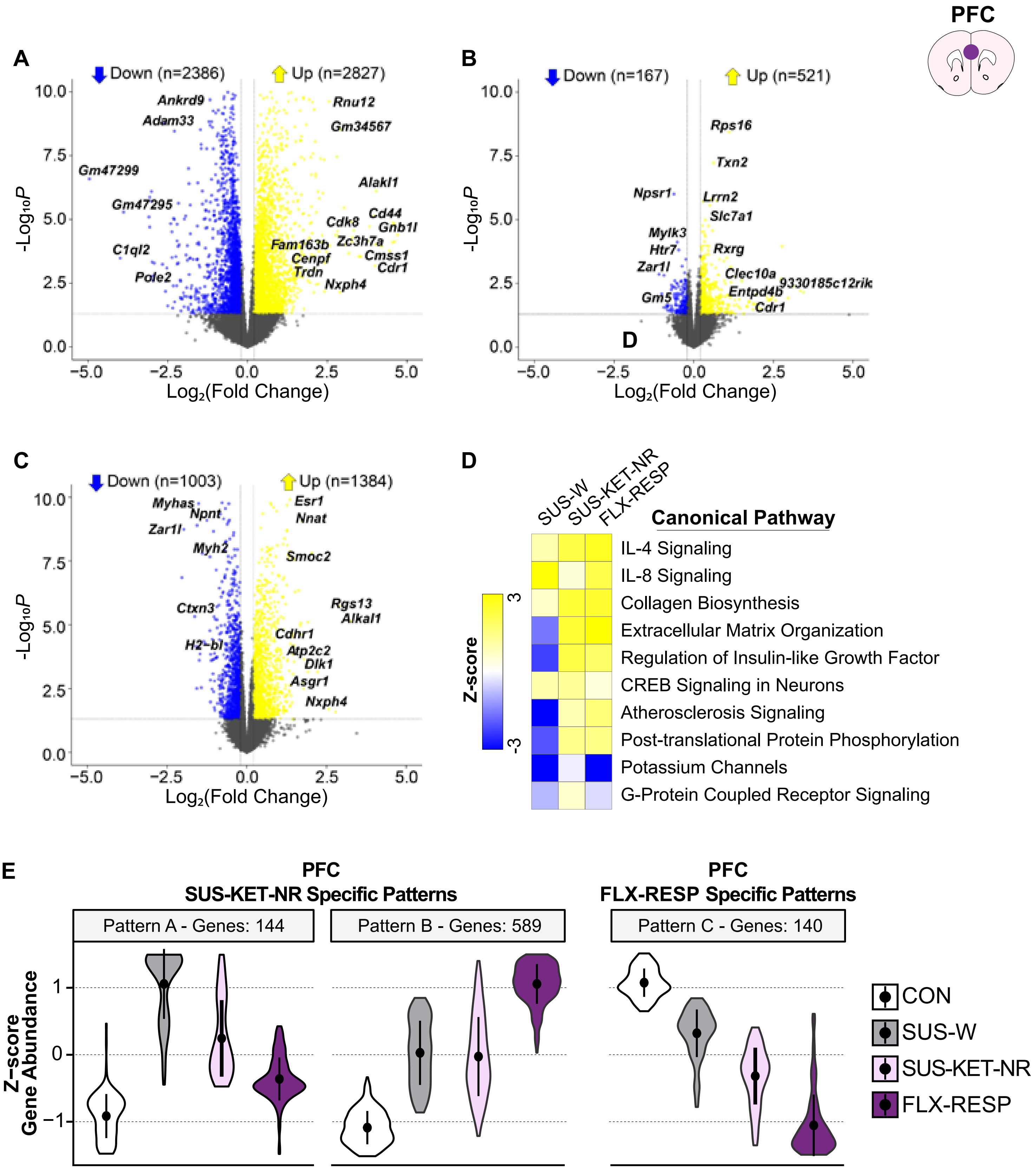

### Figure S4.

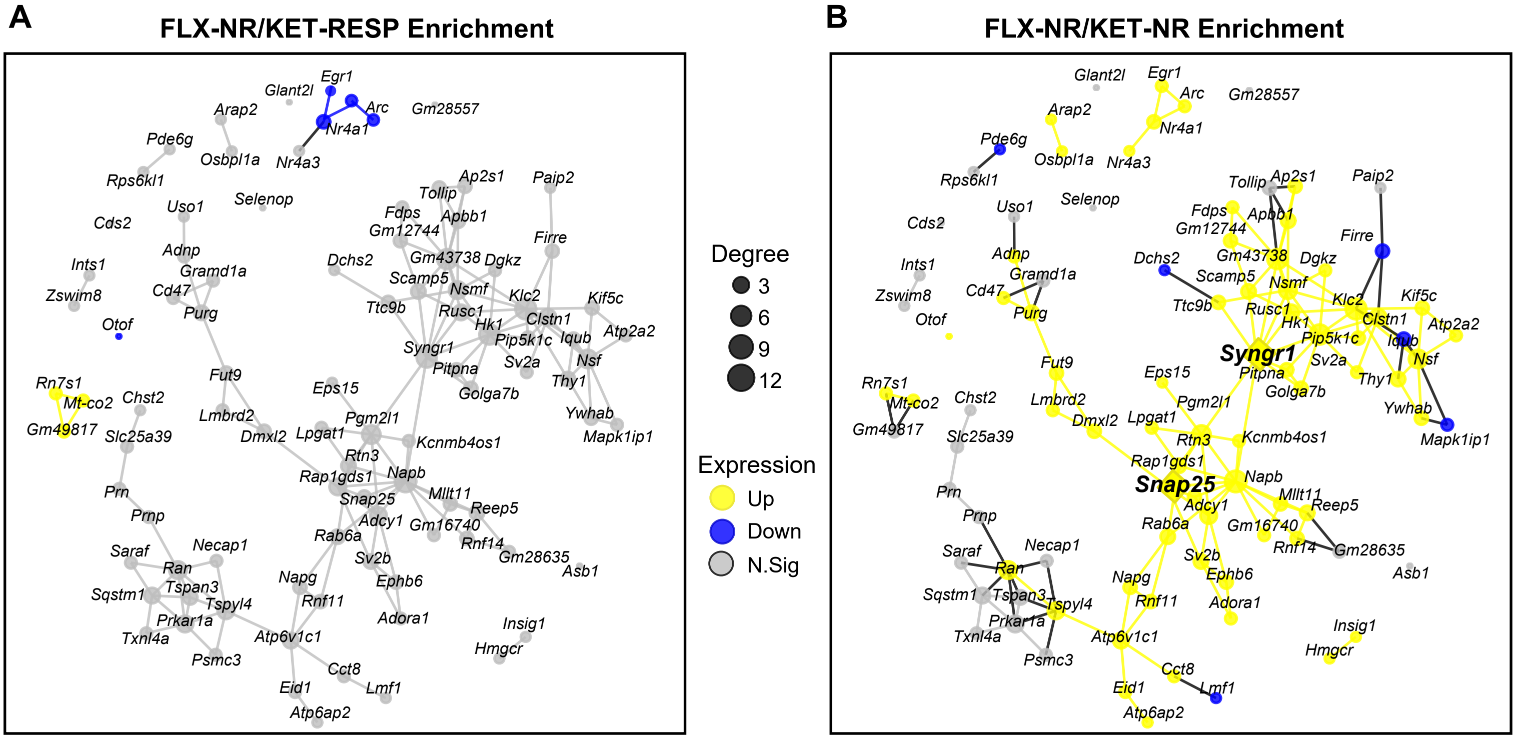
